# Cell-type-informed genotyping of mosaic focal epilepsies reveals cell-autonomous and non-cell-autonomous disease-associated transcriptional programs

**DOI:** 10.1101/2024.10.01.615793

**Authors:** Sara Bizzotto, Edward A. Stronge, Maya Talukdar, Qiwen Hu, Zinan Zhou, August Yue Huang, Brian H. Chhouk, Alissa M. D’Gama, Edward Yang, Timothy E. Green, David C. Reutens, Saul A. Mullen, Michael S. Hildebrand, Russell J Buono, Annapurna H. Poduri, Sattar Khoshkhoo, Christopher A. Walsh

## Abstract

Recent studies demonstrate growing roles for genetic mosaicism in neurodevelopmental and neuropsychiatric disorders, with the paradigm being drug-resistant pediatric focal epilepsy related to activating somatic variants in the PI3K-mTOR pathway. While identifying the genotype-associated changes at the single-cell level is fundamental to understanding disease pathophysiology, this remains technically challenging in human tissue samples with existing methods. Here, we performed single-nucleus RNA-sequencing (snRNA-seq) of 20 focal cortical dysplasia (FCD) samples removed surgically for treatment of drug-resistant epilepsy, and 10 non-FCD controls, and we developed a new approach, Genotyping Of Transcriptomes Enhanced with Nanopore sequencing (GO-TEN), that combines targeted complementary (c)DNA sequencing with snRNA-seq to perform concurrent single-nucleus genotyping and transcriptional analysis. We find that mosaic pathogenic variants in FCD do not produce a detectable novel cell identity, but instead we observe conserved cell types present both in FCD cases and non-FCD control specimens. Similarly, GO-TEN analysis shows that most pathogenic variant-carrying cells have well-differentiated neuronal or glial identities and are enriched for layer II-III excitatory neurons. We identify cell-intrinsic disruption of glutamate and GABA-A signaling pathways in variant-carrying neurons and altered intercellular signaling, making potential mechanisms for epileptogenesis in FCD. In summary, by addressing genotype-specific changes in mosaic epilepsy-associated lesions, our study highlights new potential disease mechanisms and therapeutic targets.

## Introduction

Focal cortical dysplasia (FCD) spectrum disorders represent cortical malformations that occur in the absence of other organ system involvement and are the most common cause of drug-resistant epilepsy requiring neurosurgical treatment in children^1^. FCDs are developmental brain lesions that we and others have demonstrated result from pathogenic somatic mutations that occur during cortical development, and hence are unilateral or more commonly limited to a small cortical region^2,3^. The most common FCD subtypes, FCD type 2A (FCD2A), FCD type 2B (FCD2B), and hemimegalencephaly (HME) are characterized by localized regions of abnormal cortex appearing at the histopathological level as dyslamination of the six-layer structure and the presence of rare, misoriented, dysmorphic neurons (DNs). In FCD2B and HME, another hallmark finding consists in balloon cells (BCs), which are rare, enlarged, oval-shaped, eosinophilic cells with one or more nuclei^4^. FCD type 1 (FCD1) constitutes another subtype and is defined by the presence of vertical microcolumns and/or abnormal cortical layering without DNs nor BCs^5^. A subtype of FCD is associated with oligodendroglial hyperplasia^6^. Finally, tuberous sclerosis complex (TSC) is characterized by multiple dysplastic cortical lesions, which are commonly referred to as cortical tubers, and benign sub-ependymal tumors^7^.

Pathogenic somatic variants in FCD often result in hyperactivation of PhosphatidylInositol-3 Kinase (PI3K)-mechanistic Target of Rapamycin (mTOR) pathway, which regulates cell growth, metabolism, and proliferation^8^. Gain-of-function (GoF) somatic heterozygous variants have been found in mTOR activators, such as *AKT3*, *PIK3CA*, *RHEB,* and *MTOR*^3, 9, 10^. Additionally, somatic loss-of-function (LoF) variants have been found in PI3K-mTOR pathway repressor genes such as *TSC1/2*, *DEPDC5*, *NPRL2/3,* and *PTEN.* The somatic LoF variants are usually found in the presence of a germline LoF variant in the second allele, thus causing mosaic biallelic LoF reminiscent of Knudson’s second hit model for tumor suppressors^2, 3, 11, 12^. FCD2 and HME are genetically and histopathologically very similar lesions, with the hemispheric extent and the larger fraction of variant-carrying cells in HME potentially indicating an earlier developmental stage at which the mutation occurs.

While up to 80% of dysplasias with FCD2B pathology show activating PI3K-mTOR variants^13^, how the genetic variants affect cell identities and states in the brain, and how these effects might lead to intractable epilepsy, is unclear. To address these continued unanswered questions, we performed single-nucleus RNA sequencing (snRNA-seq) of surgical specimens obtained from a cohort of 19 patients clinically diagnosed with FCD spectrum pathology and representing a variety of previously reported PI3K-mTOR pathway-associated variants^3, 13^, and compared them to 4 non-lesional cortical resections from temporal lobe epilepsy (TLE) surgery, as well as snRNA-seq data from multiple cortical brain regions in neurotypical individuals^14, 15^, including a total of 149,797 disease and 72,155 control nuclei. In addition, we also analyzed previously published data^16^ from one control and five FCD spectrum disorder cases including 2,429 and 32,345 cells, respectively. By comparing cases and controls, we show that FCD lesions do not contain any novel cell clusters other than those found in control brains at detectable frequency, nor do they display consistent shifts in cell type composition. Secondly, we developed GO-TEN, which combines 10X Genomics snRNA-seq and Oxford Nanopore (ONT) long-read sequencing of amplicons covering the variant loci as well as the unique cell barcodes, allowing genotyping of targeted loci in single nuclei while also obtaining their full transcriptional profiles. Using GO-TEN, we show that pathogenic variant-carrying cells have well differentiated cell fates and demonstrate enrichment of variants in upper layer excitatory neurons in the cortex. Furthermore, GO-TEN identified gene expression changes specific to the variant-carrying cells, as well as altered intercellular signaling. Thus, our study reveals previously unknown cell type- and genotype-specific cell-autonomous and non-cell-autonomous alterations contributing to FCD pathophysiology.

## Results

### Conserved cortical cell types between control and FCD brains

Histopathological hallmarks of FCD include the presence of DNs expressing NeuN, GFAP, excess of neurofilaments (SMI31) and phosphorylated ribosomal protein S6 (pS6), and in certain subtypes of eosinophilic BCs^4^. In order to determine the cell types and identities present in FCD lesions and whether DNs and BCs are transcriptionally distinguishable, we performed snRNA-seq of 20 fresh-frozen surgically resected brain tissues for a total of 19 FCD spectrum disorder cases, including ten FCD2, seven HME (two samples in the lesion were obtained from the same individual), one FCD1A, and one FCD not otherwise specified (NOS), and compared them with control snRNA-seq data from six non-FCD brains (Fig. 1a, Table 1, Extended Data Table 1, Extended Data Fig. 1a and Methods). The non-FCD controls consisted of anterior temporal neocortex surgical resections from four patients with TLE as non-lesional epilepsy controls (Brodmann area 38), as well as eleven prefrontal cortex (Brodmann area BA9) and occipital cortex samples (primary and secondary visual cortices, BA17 and BA18) from two neurotypical post-mortem brains^14, 15^. Additionally, we analyzed snRNA-seq data obtained from cortical tissue previously published by Chung et al.^16^ consisting of one control and five FCD cases (including two FCD2/HME, two TSC, and one FCD-NOS). To control for potential technical biases, we performed the analyses first including only data generated in our lab, and then again additionally including data published by Chung et al.^16^. Pathogenic variants were previously identified^3, 13^ in 16 FCD patients in our cohort, including one *TSC1*, four *PIK3CA* and eight *MTOR* somatic variants, one somatic copy number gain in the long (q) arm of chromosome 1 including *AKT3*, one *SLC35A2* somatic variant, and one *DEPDC5* germline variant. Somatic variants had bulk VAFs spanning 2.3% to 25.8% (Table 1). Overall, a total of 149,797 single cells from FCD cases and 72,155 control cells passed quality control filters (Extended Data Fig. 1b, and Methods).

**Figure 1.**
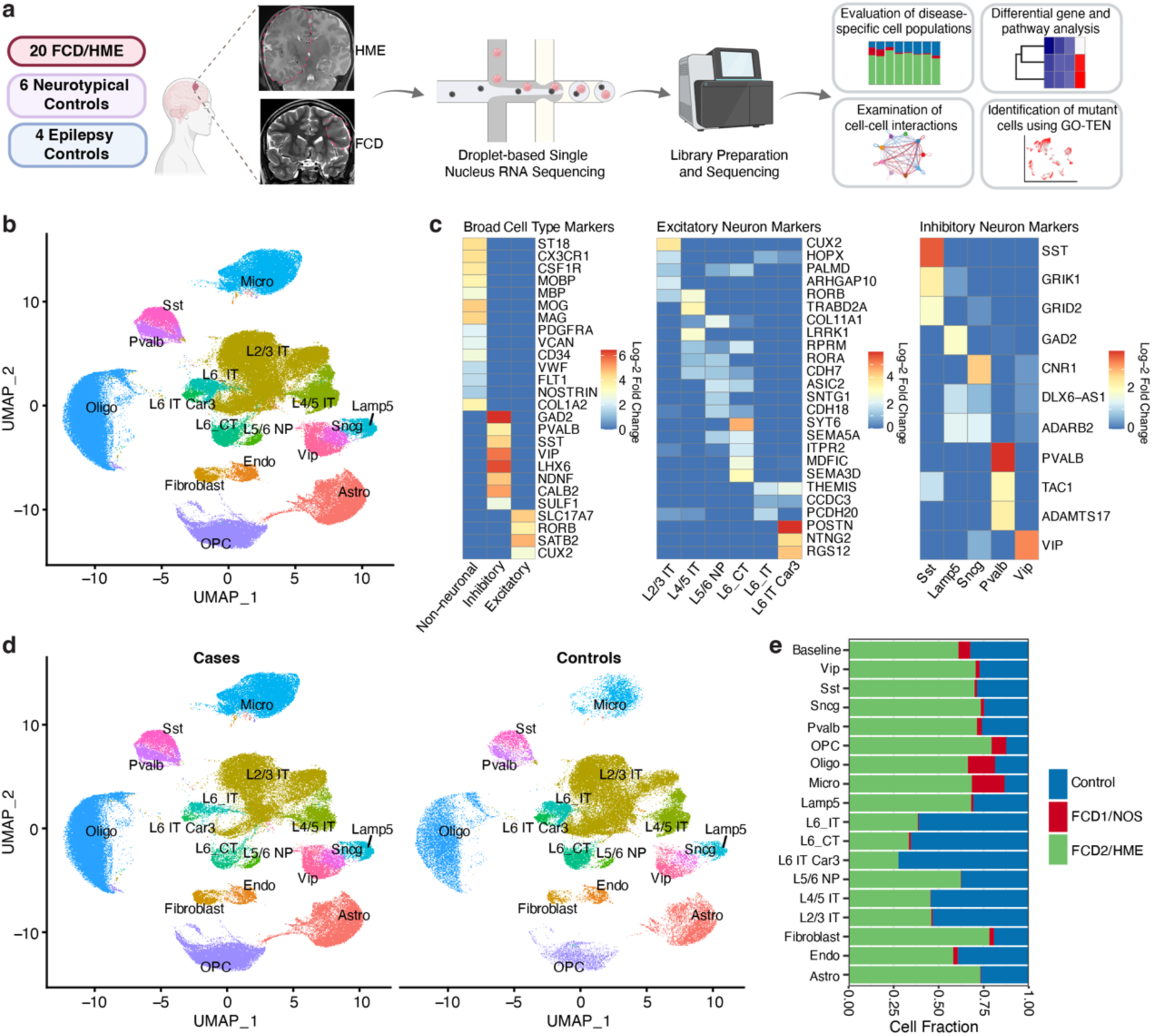
Conserved cortical cell types between control and FCD brains. (**a**) Schematic of the experimental and analytical workflow. Representative coronal T2 MRI images for HME (E174) and FCD2B (E364) demarcate the abnormality with dashed lines. (**b**) Uniform Manifold Approximation and Projection (UMAP) showing cell type clusters identified in the integrated disease and control samples. (**c**) Heatmaps showing the expression of known cell type markers used for annotation. (**d**) UMAPs showing conserved cell type clusters between cases and controls. (**e**) Contribution of each disease and control group to cell types. Differential cell type composition analysis was done using a one-way ANOVA test, which showed non-significant shifts.

**Table 1.** Sample cohort.

| Table 1. Sample cohort |  |  |  |  |  |  |
| --- | --- | --- | --- | --- | --- | --- |
| Sample ID | Case ID | Brain region | Sample type | Diagnosis | Variant | VAF (%) |
| E274 | E274 | L-CC | FF surgical | HME | *PIK3CA*<br>p.E542K<sup>13</sup> | 25.8 |
| E241 | E241 | L-CC | FF surgical | HME | *PIK3CA*<br>p.E545K<sup>13</sup> | 23.5 |
| E348 | E348 | R-CC | FF surgical | HME | *PIK3CA*<br>p.E545K<sup>13</sup> | 16.4 |
| E174-1 | E174 | R-FTC | FF surgical | HME | *PIK3CA*<br>p.E542K<sup>3</sup> | 15.9-17.4 |
| E174-2 | E174 | L-CC | FF surgical | HME | *PIK3CA*<br>p.E542K | 13.9 |
| E254 | E254 | R-CC | FF surgical | HME | *MTOR*<br>p.C1483R<sup>13</sup> | 23.7 |
| E276 | E276 | - | FF surgical | HME | *MTOR*<br>p.C1483Y<sup>13</sup> | 11.7 |
| FC5801 | FC5801 | L-H | FF surgical | FCD2B | *MTOR*<br>p.C1483R<sup>3</sup> | 10.0-10.6 |
| E286 | E286 | R-CC | FF surgical | FCD2B | *MTOR*<br>p.S2215F<sup>13</sup> | 5.8 |
| E364 | E364 | L-FC | FF surgical | FCD2 | *MTOR*<br>p.T1977K<sup>13</sup> | 4.7 |
| E244 | E244 | L-FC | FF surgical | FCD2 | *MTOR*<br>p.T1977R<sup>3</sup> | 2.8-4.7 |
| E316 | E316 | R-POTC | FF surgical | FCD2 | *MTOR*<br>p.S2215Y<sup>13</sup> | 4.5 |
| FC5501 | FC5501 | L-FPC | FF surgical | FCD2B | *MTOR*<br>p.L1460P<sup>3</sup> | 2.3-2.6 |
| FC6401 | FC6401 | L-FC | FF surgical | FCD2B | *TSC1*<br>p.Q55X<sup>3</sup> | 5.1-6.7 |
| E269 | E269 | R-FTC | FF surgical | FCD-NOS | *SLC35A2*<br>p.M1I<sup>13</sup> | 3.0 |
| E203 | E203 | L-FTC | FF surgical | FCD2A | 1q dup<br>(*AKT3*)<sup>13</sup> | NA |
| FC10801 | FC10801 | L-FC | FF surgical | FCD2B | *DEPDC5*<br>p.G204fs | germline |
| HE3601 | HE3601 | R-CC | FF surgical | HME | - | - |
| FC5901 | FC5901 | L-FC | FF surgical | FCD2B | - | - |
| E216 | E216 | L-FTC | FF surgical | FCD1A | - | - |
| UMB4638-S1 | UMB4638 | L-PFC | FF<br>postmortem | NT ctrl | - | - |
| UMB4638-S2 | UMB4638 | L-OC-V1 | FF<br>postmortem | NT ctrl | - | - |
| UMB4638-S3 | UMB4638 | L-OC-V1 | FF<br>postmortem | NT ctrl | - | - |
| UMB4638-S4 | UMB4638 | L-OC-V1 | FF<br>postmortem | NT ctrl | - | - |
| UMB4638-S5 | UMB4638 | L-OC-V2 | FF<br>postmortem | NT ctrl | - | - |
| UMB4638-S6 | UMB4638 | L-OC-V2 | FF<br>postmortem | NT ctrl | - | - |
| UMB4643-S1 | UMB4643 | L-PFC | FF<br>postmortem | NT ctrl | - | - |
| UMB4643-S2 | UMB4643 | L-OC-V1 | FF<br>postmortem | NT ctrl | - | - |
| UMB4643-S3 | UMB4643 | L-OC-V1 | FF<br>postmortem | NT ctrl | - | - |
| UMB4643-S4 | UMB4643 | L-OC-V2 | FF<br>postmortem | NT ctrl | - | - |
| UMB4643-S5 | UMB4643 | L-OC-V2 | FF<br>postmortem | NT ctrl | - | - |
| TLE36 | TLE36 | TC | FF surgical | TLE ctrl | - | - |
| TLE6 | TLE6 | TC | FF surgical | TLE ctrl | - | - |
| TLE7 | TLE7 | TC | FF surgical | TLE ctrl | - | - |
| FC10101 | FC10101 | TC | FF surgical | TLE ctrl | - | - |

Abbreviations: R-FTC, right frontotemporal cortex; L-CC, left cerebral cortex; R-CC, right cerebral cortex; L-FPC, left frontoparietal cortex; L-H, left hemisphere; L-FC, left frontal cortex; R-POTC, right parieto-occipital temporal cortex; L-FTC, left frontotemporal cortex; L-PFC, left prefrontal cortex; L-OC-V1, left occipital cortex primary visual; L-OC-V2, left occipital cortex secondary visual; TC, temporal cortex; FF, fresh-frozen; NT, neurotypical; TLE, temporal lobe epilepsy

We processed and integrated all case and control samples, controlled for ambient RNA contamination^17^, and performed cell type annotation using canonical brain cell type markers^18^ (Fig. 1b, c, Extended Data Fig. 2, Extended Data Fig. 3a-c and Methods). We identified the most common known cortical cell types including six excitatory neuron subtypes, five inhibitory neuron subtypes, oligodendrocyte precursor cells (OPCs), mature oligodendrocytes, astrocytes and microglia, as well as fibroblasts and endothelial cells. Excitatory neuron subtypes included layer (L) 2/3, L4/5, L6 and L6-Car3 intratelencephalic (IT) neurons, L5/6 near-projecting (NP) neurons, and L6 cortico-thalamic (CT) neurons. Inhibitory neuron subtypes included Sst, Lamp5, Sncg, Pvalb, and Vip. We found that all clusters in the dataset contained cells from both cases and controls. Furthermore, we could not identify distinguishable new clusters specific to FCD cases and representing cells of an abnormal or novel identity potentially representing DNs and/or BCs (Fig. 1d), despite apparent histological abnormalities in these hallmark, but rare, cell types. Several possible explanations for this include 1] these cell types are rare, and so may require very large samples to be captured, 2] BCs are frequently multinucleated and hence may be filtered by QC doublet removal methods, or 3] they do not represent transcriptionally distinct cellular identities in the dysplastic cortex.

We then looked for possible cell type composition shifts in FCD brains by comparing the contribution of FCD2/HME, FCD1/FCD-NOS, and controls to cell types, and found no differences that reached statistical significance (ANOVA test; Fig. 1e and Methods). The same results were replicated when including data from Chung et al (Extended Data Fig. 3d).

Thus, overall our analysis shows no evidence of new transcriptionally distinguishable abnormal cell clusters in FCD, suggesting that the morphological alterations that characterize DNs and BCs in FCD tissue are not translated into a new cell identity as determined by gene expression, or that such changes are sufficiently modest that they will require larger sample sizes to capture. Furthermore, we found no consistent cell type proportion shifts between cases and controls, indicating that FCD dysplastic cortex does not present major alterations in cellular composition compared to the normal non-dysplastic cortex.

### FCD brains show altered energy metabolism and microglial immune activation

Having identified the major cell types in FCD, differential gene expression (DGE) analysis between cases (including all FCD subtypes) versus controls (Methods) identified multiple differentially regulated transcriptional programs. Gene set enrichment analysis (GSEA) utilizing the fifty Hallmark Pathways^19, 20^ showed down-regulation of oxidative phosphorylation in all cell types in the FCD group. Almost all cell types also showed downregulation of MYC targets and KRAS signaling, and several, including excitatory and inhibitory neurons sub-types, showed downregulation of hypoxia and reactive oxygen species pathways, as well as glycolysis, fatty acid metabolism, protein secretion, adipogenesis, and mTORC1 and PI3K-AKT-MTOR signaling (Fig. 2a). These results suggest that FCD brains are affected by global alterations in energy metabolism and biosynthesis, which may significantly alter key cell features such as axonal transport and synaptic function^21^. The paradoxical downregulation of pathways associated with cell growth and proliferation, such as KRAS, MYC and PI3K-AKT-MTOR may reflect a compensatory mechanism due to PI3K-mTOR hyperactivity in variant-carrying cells (see below).

**Figure 2.**
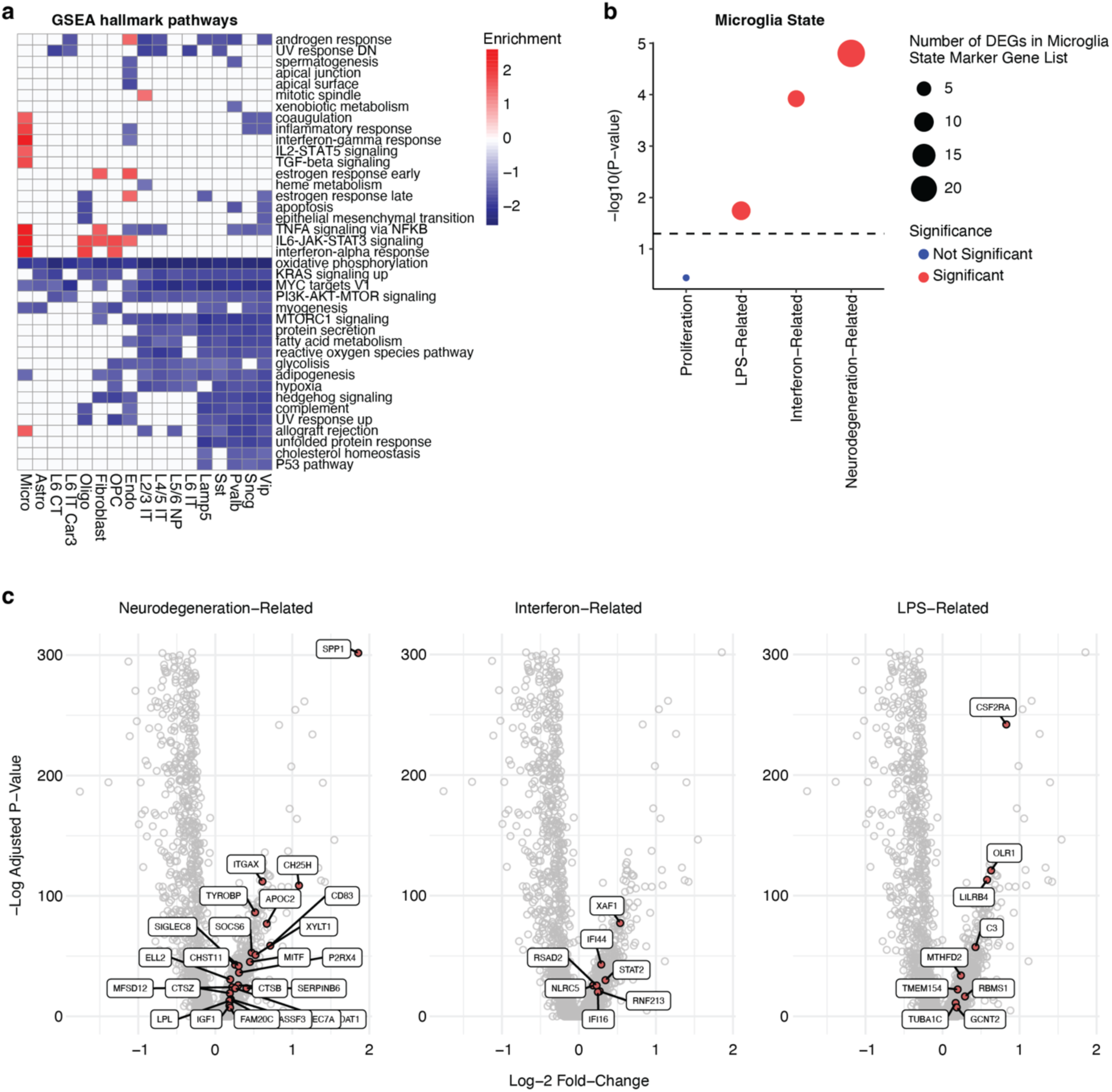
Alteration of energy metabolism and microglia immune activation in FCD. (**a**) Heatmap showing cell-type-associated pathways positively (red) and negatively (blue) enriched in cases compared to controls obtained through Gene Set Enrichment Analysis (GSEA). (**b**) Number of differentially expressed genes (DEGs) in microglia state marker gene list, highlighting microglia activation in disease. *P*-values were calculated using a Friedman test. Significance for adj.*p* < 0.05. (**c**) Differentially expressed genes in microglia state marker gene lists related to neurodegeneration, interferon and lipopolysaccharide (LPS).

GSEA also revealed FCD-associated immune activation in microglia, where we found upregulation of terms such as inflammatory response, interferon-gamma response, IL2-STAT5 signaling, TGF-beta signaling, TNFA signaling via NFKB, IL6-JAK-STAT3 signaling and interferon-alpha response (Fig. 2a). To further dissect the alterations of microglial gene expression we observed in FCD, we utilized a previously published single-cell (sc)RNA-seq atlas of various microglial states derived from both human and mouse brain tissue^22^. We identified a significant overlap between differentially expressed genes (DEGs) that were upregulated in FCD microglia and microglial states associated with LPS-induced immune activation, interferon response, and neurodegeneration (Fig. 2b, c). Taken together, our data highlight a role for microglia activation in FCD pathology that may be a cause or a consequence of epileptic seizures and mirrors microglial activation seen in other neurological disorders^23^.

### Altered signaling between neuronal subtypes in FCD

In order to identify mechanisms potentially contributing to epileptic seizures in FCD patients, we performed GSEA by specifically interrogating the synaptic gene ontology (SynGO) database^24^, as well as lists of genes associated with epilepsy based on Macnee et al.^25^, DisGeNET (https://www.disgenet.com)^26^, and the Simons Foundation Autism Research Initiative (SFARI) database (https://gene.sfari.org) (see Methods). The latter was used due to epilepsy being a known co-morbidity for autism spectrum disorder (ASD). We found that several synapse-related terms showed enrichment or depletion in cases (Fig. 3a). All cell types showed downregulation of gene sets related to pre- and post-synaptic translation. In excitatory neurons specifically, we found upregulation of terms such as integral component of postsynaptic density membrane, integral component of presynaptic membrane, modulation of chemical synaptic transmission, and regulation of postsynaptic membrane neurotransmitter receptor levels. Furthermore, we saw global downregulation of Macnee et al. and DisGeNET^26^ epilepsy-associated genes. Finally, we found upregulation of SFARI genes specifically in excitatory neurons (Fig. 3a). Synaptic dysfunction has an established role in epilepsy^27^. Our GSEA analysis performed on epilepsy-related pathways revealed a general disruption of synaptic translation in FCD tissue, while regulation of pre- and post-synaptic membrane, synaptic transmission and neurotransmitter levels seemed affected specifically in FCD excitatory neurons.

**Figure 3.**
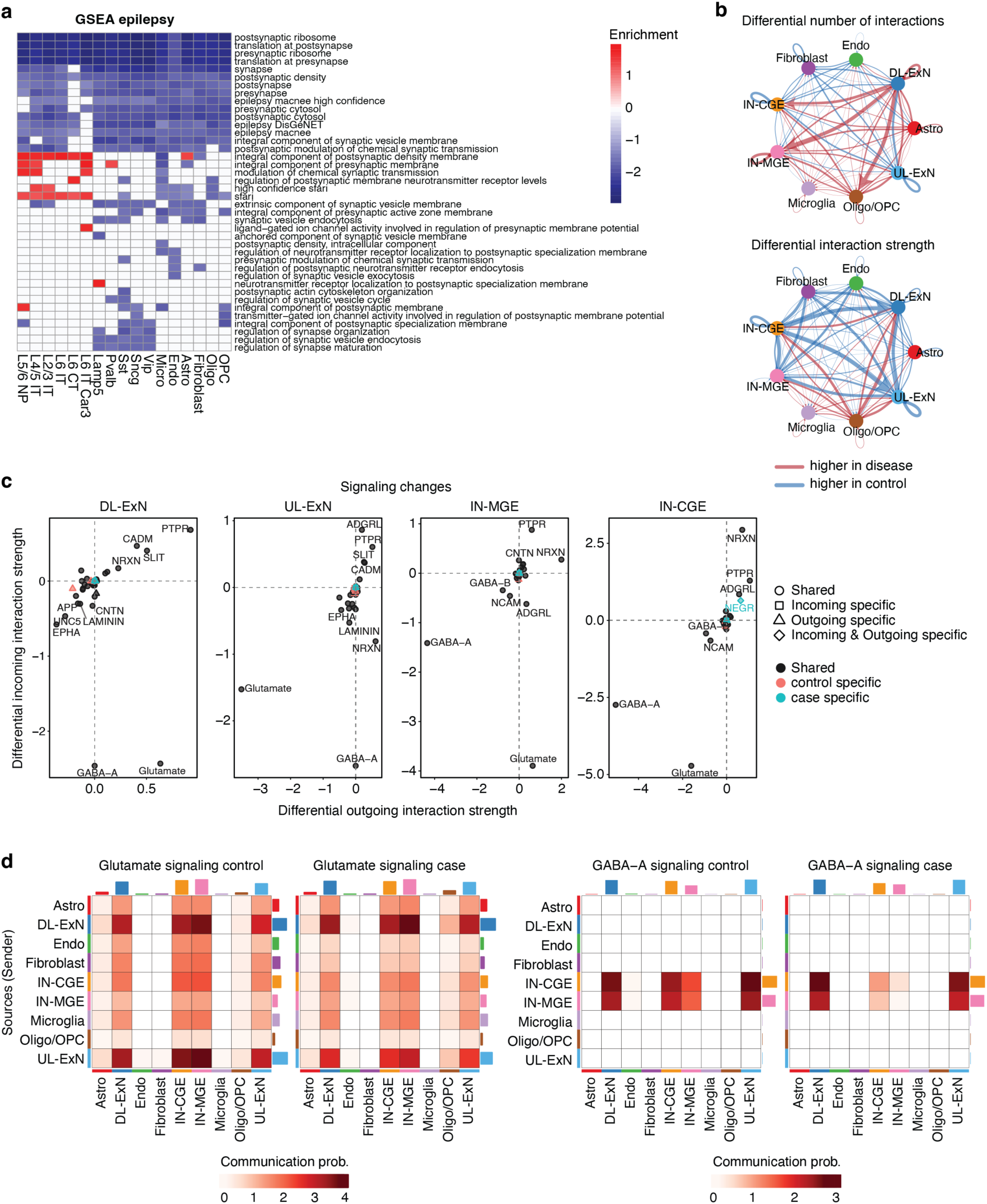
Altered signaling between neuronal subtypes in FCD. (**a**) Heatmap showing cell-type-associated pathways positively (red) and negatively (blue) enriched in cases compared to controls, obtained through GSEA run on epilepsy-related pathways. (**b**) Circle plot where edge widths represent the case versus control differential number of interactions (top panel) and differential interaction strength (bottom panel) between cell groups calculated with CellChat. (**c**) Differential incoming and outgoing signaling changes for neuronal cell groups calculated with CellChat. Positive and negative values indicate respectively increased and decreased signaling in disease compared to control. (**d**) Glutamate and GABA-A communication probability between cell groups calculated with CellChat in cases and controls. The signaling sender (outgoing) cell group is reported in the y-axis, while the signaling receiver (incoming) is reported in the x-axis.

To further explore changes in intercellular signaling that could explain FCD-related seizure susceptibility, we applied CellChat, a tool that quantitatively infers intercellular communication networks from scRNA-seq data based on the expression of known receptor/ligand pairs^28^ (Methods). To maximize statistical power for this analysis, we divided neurons into four major groups: deep-layer excitatory neurons (DL-ExN, including L4/5 IT, L5/6 NP, L6 CT, L6 IT and L6 IT Car3), upper-layer excitatory neurons (UL-ExN, including only L2/3 IT), caudal ganglionic eminence (CGE)-derived interneurons (IN-CGE, made of Vip, Sncg and Lamp5), and medial ganglionic eminence (MGE)-derived interneurons (IN-MGE, including Sst and Pvalb)^29^. We found an increased number of inferred interactions going from DL-ExN to IN-CGE and IN-MGE, as well as to oligodendrocytes and OPCs (Fig. 3b), however, the strength of interactions originating from IN-CGE and towards DL-ExN and UL-ExN appeared to be weaker (Fig. 3b). This pointed towards a specific perturbation of intercellular signaling potentially involving increased excitation from DL-ExN and decreased inhibition of excitatory neurons by IN-CGE. We further explored the specific features driving these interactions, and identified changes in glutamate and GABA-A signaling in all four neuronal subgroups. More precisely, we found increased outgoing glutamate signaling from DL-ExN and a decreased outgoing glutamate signaling from UL-ExN. Outgoing GABA-A signaling was reduced in both interneuron subgroups, and incoming GABA-A signaling was reduced in all neurons (Fig. 3c, d). These results suggest specific directional changes in synaptic connectivity and function involving excitatory and inhibitory neurons in FCD that could account for circuit dysfunction and epileptogenesis.

### Cell type-informed genotyping of pathogenic somatic mutations

Identifying cell states associated with the pathogenic genotype in mosaic neurological conditions represents a technological hurdle as it requires sequencing both the DNA and RNA from single cells^30, 31^. Although recent technologies such as G&T-seq (Genome and Transcriptome sequencing)^32^ and ResolveOME^33^ have enabled such studies, these methods remain low-throughput and cost prohibitive. Consequently, we developed Genotyping Of Transcriptomes Enhanced with Nanopore sequencing (GO-TEN) which represents a modification of Genotyping of Transcriptomes (GoT)^34^ and combines 10X snRNA-seq with an orthogonal genotyping strategy based on ONT long-read sequencing. GO-TEN can be applied to the full-length cDNA that did not undergo fragmentation and library preparation, obtained from an intermediate step in the 10X Genomics snRNA-seq protocol, and entails generation of on-target amplicons that capture the cell barcodes and unique molecular identifiers (UMIs) for the transcripts, as well as the genomic locus corresponding to the variant of interest. This step is performed using biotinylated primers that allow pull-down enrichment of the amplified transcripts. A hemi-nested PCR is then performed to further improve specificity before library preparation and long-read sequencing. At the end of the protocol, the barcode information is used to map genotyped cells to the Illumina short-read snRNA-seq data obtained from the same cDNA pool. To improve genotyping accuracy, we used a custom adaption of the Bayesian genotyper from MosaicHunter optimized for snRNA-seq data^35, 36^ (Fig. 4a and Methods).

**Figure 4.**
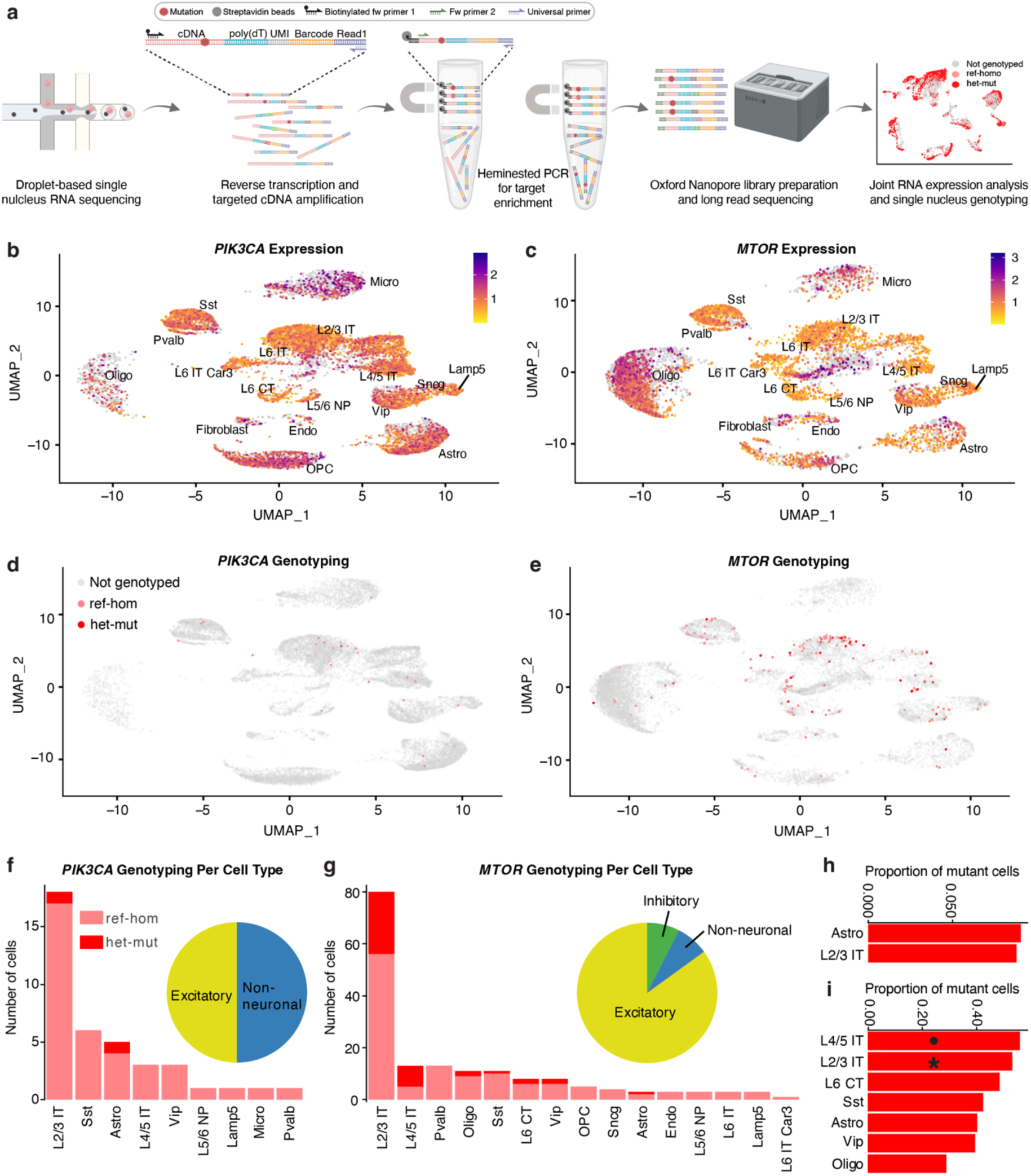
Cell type-informed genotyping of pathogenic somatic variants. (**a**) Schematic of the GO-TEN experimental workflow. (**b-c**) *PIK3CA* and *MTOR* expression across cell types based on snRNA-seq. (**d-e**) UMAPs showing *PIK3CA* and *MTOR* reference homozygous and heterozygous mutant cells genotyped with GO-TEN high-specificity approach. (**f**-**g**) Number of cells genotyped with GO-TEN high-specificity approach as reference homozygous and heterozygous mutant. Pie charts show the contribution of broad cell categories to mutant cells. (**h**-**i**) Number of heterozygous mutant cells for *PIK3CA* and *MTOR* normalized for the total number of genotyped cells for each mutant cell type. Data also include cells genotyped using the high-sensitivity approach. Asterisk (adj. p<0.05) and black circle (p<0.05) indicate significant enrichment of the mutation for a particular cell type (Fisher test).

We used GO-TEN to perform single-cell genotyping of surgical samples from three patients in our cohort carrying known recurrent activating missense PI3K-mTOR pathway variants. The first patient (E174) was clinically diagnosed with HME and carried a *PIK3CA* p.E542K variant with ∼28% mosaicism (bulk VAF = 13.9 – 17.4%)^3^. The second patient (FC5501) was diagnosed with FCD2B, and carried an *MTOR* p.L1460P variant with ∼5% mosaicism (bulk VAF = 2.3 – 2.6%)^3^. The third patient (FC5801) was also diagnosed with FCD2B, and carried an *MTOR* p.C1483R variant with ∼20% mosaicism (bulk VAF = 10.0-10.6%)^3^. We performed genotype analysis using two methods: 1) a more stringent approach to maximize specificity, and 2) a less stringent approach to maximize sensitivity (see Methods). With the high-sensitivity approach, we genotyped a total of 5500 nuclei across all three cases. Of these, 3682 were homozygous for the reference allele (ref-hom) and 1818 were heterozygous for the mutant allele (het-mut). 1740 cells were genotyped for the *PIK3CA* locus, of which 1607 were ref-hom, and 133 (7.6%) were het-mut. *MTOR* loci were genotyped in a total of 3760 cells (3114 for FC5501 and 646 for FC5801), of which 2075 were ref-hom, and 1685 (44.8%) were het-mut. The fraction of het-mut cells in each *MTOR* case were 43.4% and 51.4% for FC5501 and FC5801, respectively. We sought to understand factors that affect genotyping efficiency by analyzing several covariates. We found that the expression level of a gene in a given cell positively correlated with the probability of genotyping, which is expected since GO-TEN genotypes RNA molecules. Surprisingly however, we also found that some cell types had increased or decreased probability of being genotyped for a specific gene mutation (Extended Data Fig. 4a). This represents an area of future investigation; however, we speculate that it may be a consequence of the expression of specific transcript isoforms that may be more or less efficiently captured by GO-TEN.

GO-TEN allowed us to examine the distribution of mutant cells across cell types in our dataset (Fig. 4b-g, Extended Data Fig. 4b-i). In order to identify the variant-carrying cell types with high confidence, we considered only nuclei genotyped with the high-specificity approach (Fig. 4d-g). We found that the *PIK3CA* variant was distributed evenly between excitatory neurons and glial cells: more precisely, L2/3 IT excitatory neurons and astrocytes (Fig. 4f). The *MTOR* variants, on the other hand, were found primarily in excitatory neurons, but mutant cells were also identified in some inhibitory neurons and glial cells. Indeed, mutant cells for *MTOR* were identified in L2/3 IT, L4/5 IT, L6 CT excitatory neurons, and Sst and Vip interneurons, as well as oligodendrocytes and astrocytes (Fig. 4g) consistent with the mutation arising in a neural epithelial progenitor cell. As expected, variant carrying cells were rarely pericytes or microglia at levels consistent with doublet rates, aligning with the fact that these cells have a embryological divergent origin. The shared presence of the variants in both excitatory and inhibitory neurons could be interpreted as suggesting an early origin of the mutation, but more recent studies^15, 37–39^ have suggested a shared origin of these neuronal types even relatively late in neurogenesis. Thus, we interpret the cells containing the variant as most likely consistent with the normal neural lineages, rather than strongly suggesting a lineage that is altered by the variants.

We next tested for enrichment of FCD pathogenic mutations in specific cell types. In order to increase our statistical power, we implemented cell numbers by considering also cells genotyped with the high-sensitivity approach. Since some cell types are positively associated with a higher probability of genotyping (Extended Data Fig. 4a), we tested for enrichment by normalizing the number of het-mut cells on genotyped cells. The *PIK3CA* variant was only identified in L2/3 IT neurons or astrocytes (Fig. 4h), although the limited sample size prevented this enrichment from being statistically significant. Enrichment of *MTOR* variants on the other hand was statistically significant in L2/3 IT neurons (adjusted *p*-value < 0.05; Fig. 4i), and to a lower extent in L4/5 IT neurons (nominal *p*-value < 0.05, Fig. 4i). Thus, our cell-type-informed genotyping indicates that FCD pathogenic mutations, while detectable across several cell types, show enrichment in middle-to-upper-layer excitatory neurons. This upper layer enrichment likely reflects at least in part the developmental timing of mutation occurrence, since it mirrors upper layer excitatory neuron enrichment of functionally neutral somatic variants at low VAF in normal brain^15, 35^. However, such enrichment could also reflect positive or negative selection affecting the variant-carrying clones, or shifts in the progenitor potential induced by the variant. In any event, this enrichment implicates excitatory neurons with layer 2/3 transcriptional identity in the pathogenesis of epilepsy in FCD.

### Genotype-associated transcriptional alterations

GO-TEN additionally allowed us to explore DEGs and pathways between ref-hom and het-mut cells in our dataset. To increase statistical power, we used data from nuclei genotyped with the high-sensitivity approach; however, we restricted the analyses to the cell types carrying the variant in the high-specificity dataset. GSEA using the Hallmark Pathways gave very similar results to the non-genotype-informed disease versus control analysis. Consistent with that, in all mutated cell types we found downregulation of oxidative phosphorylation, KRAS signaling and MYC targets V1. Among the upregulated pathways specifically in mutant astrocytes and oligodendrocytes, we found IL6-JAK-STAT3 signaling and interferon-alpha response, indicating activation of inflammatory signaling. We also found downregulation of PI3K-AKT-MTOR and MTORC1 signaling in the variant-carrying neurons, likely a compensatory response to variant-induced mTOR hyperactivity. Moreover, we again detected downregulation of protein secretion, TNFA signaling via NFKB, glycolysis, adipogenesis, fatty acid metabolism, hypoxia and reactive oxygen species pathway. One Hallmark Pathway was found upregulated in L2/3 IT neurons: namely, mitotic spindle (Fig. 5a). Although neurons are post-mitotic cells, mitotic spindle regulators include cytoskeleton-associated genes that are important also for neuronal migration, molecule transport, and neuron projection development. Indeed, the leading-edge genes driving this association included *CLIP1* and *CLIP2*, *KIF1B*, and several myosin and filamin genes.

**Figure 5.**
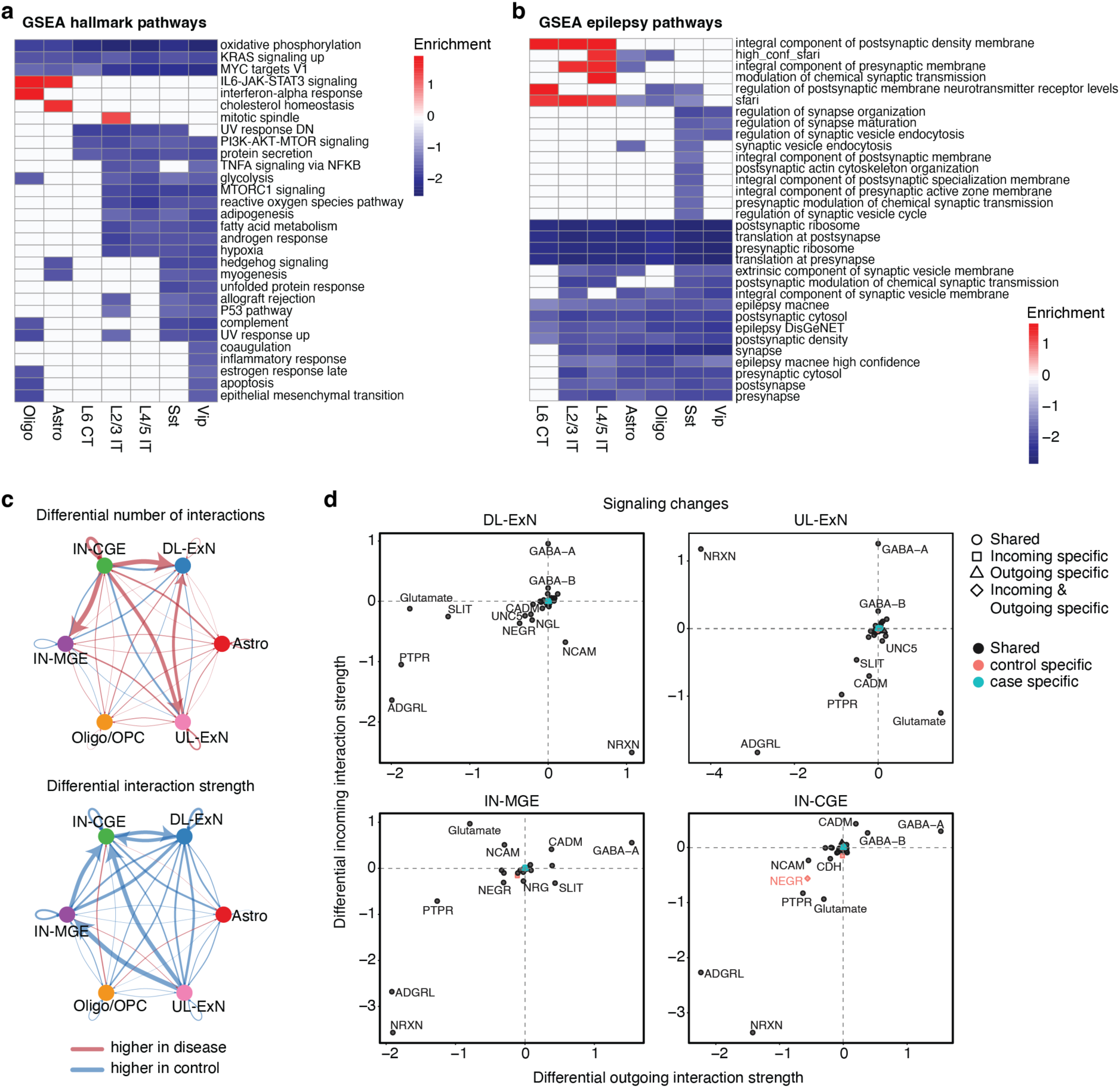
Genotype-associated transcriptional alterations. (**a**) Heatmap showing cell-type-associated GSEA pathways positively (red) and negatively (blue) enriched in heterozygous mutant compared to reference homozygous cells. The analysis was done by pooling all cells genotyped with GO-TEN from different cases. (**b**) Epilepsy-associated pathways GSEA enriched for each cell type in heterozygous mutant compared to reference homozygous cells. (**c**) Circle plot where edge widths represent the heterozygous mutant versus reference homozygous differential number of interactions (top panel) and differential interaction strength (bottom panel) between cell groups calculated with CellChat. (**d**) Differential incoming and outgoing signaling changes for neuronal cell groups calculated with CellChat. Positive and negative values indicate respectively increased and decreased signaling in heterozygous mutant compared to reference homozygous cells.

We next interrogated epilepsy-associated pathways including SynGO, Macnee et al., DisGeNET and SFARI. Again, results were similar to those obtained for the non-genotype-informed disease versus control comparison. We found upregulation of the integral component of postsynaptic density membrane and of SFARI genes in excitatory neurons specifically. Upregulation of postsynaptic membrane neurotransmitter receptor levels was found in L6 CT neurons, modulation of chemical synaptic transmission was upregulated in L4/5 IT neurons, and integral component of presynaptic membrane was upregulated in L2/3 IT and L4/5 IT neurons. Finally, we found downregulation of translation at both pre- and post-synapse and of epilepsy-associated genes (both Macnee and DisGeNET) across all mutant cell types (Fig. 5b). The striking similarities between mutant vs wild-type and FCD vs control DGEA suggests that the pathogenic variants in FCD cause disease through both cell-autonomous and non-cell-autonomous mechanisms.

We next compared het-mut nuclei to ref-homo nuclei using CellChat, which revealed increased number of interactions from variant-carrying IN-CGE to IN-MGE, DL-ExN, and UL-ExN, as well as from UL-ExN to DL-ExN (Fig. 5c). However, the strength of interactions between different interneuronal and excitatory neuron clusters was lower overall (Fig. 5c). Incoming and outgoing signaling changes in neurons highlighted glutamate and GABA-A signaling. However, a stronger effect was detected for neurexin (NRXN), receptor-type protein tyrosine phosphatase (PTPR), and adhesion G protein-coupled receptor L (ADGRL) signaling (Fig. 5d), which points towards disruptions in synaptic organization and signal transduction in variant-carrying neurons. Thus, by comparing gene expression changes in mutant and wild-type cells, our data identified global dysfunction in intercellular communication that affected all neurons irrespective of their genotype.

## Discussion

In this study, using high-throughput single-cell transcriptomics of mosaic FCD spectrum brain lesions harboring somatic activating PI3K-mTOR variants, we showed that mutant cells have preserved cell identities comparable to controls. While this finding may appear contradictory to the findings of a previously published study^16^, joint analysis of our data and the raw data from Chung et al, once corrected for ambient RNA contamination^17^, was consistent in that it did not identify any transcriptionally distinct abnormal cell clusters in FCD. Furthermore, we observed that all cell clusters were conserved between the two groups, and disease variant-carrying cells were almost invariably well-differentiated neuronal and glial cell types. Thus, somatic variants in FCD do not appear to create new, disease-associated cell identities detectable at this sample size. Although we must acknowledge that sequencing more cells per patient could allow to identify very rare cell states present in the tissue, our results favor the idea that DNs and BCs, in spite of being histological hallmarks of FCD pathology, may be very rare, filtered out by doublet screening, or have transcriptomes that overlap other cell types. This is also consistent with prior reports of normal-appearing neurons in FCD also carrying pathogenic somatic variants^2^.

We found no significant change in cell type proportions between cases and controls. Although this analysis may be limited by the non-homogeneous sampling of cortical tissue across all cases and controls, these results still indicate that the FCD-associated dysplastic cortex has a cell type composition similar to non-dysplastic brains. The enrichment of *MTOR* variants in middle-to-upper layer excitatory neurons is quite consistent with enrichment of late-occurring, functionally neutral variants in these same cell types^15, 35^, further suggesting that the major impact of *MTOR* variants is not creating new cell identifies, but instead altering the transcriptomic state of well differentiated cells. Interestingly, co-occurrence of somatic variants in excitatory and inhibitory neurons confirms prior reports of variant-carrying interneurons in FCD^40^, and is consistent with the claim of a shared dorsal progenitor for these two neuronal types^15, 38^.

Our DGE and GSE analyses identified global alterations in metabolism, biosynthesis and growth-related pathways common to multiple cell types in FCD. This suggests the presence of both cell-autonomous and non-cell-autonomous metabolic changes affecting FCD tissue. In addition, we found evidence of microglial immune activation that may either be a contributing factor or a consequence of the epileptic seizures affecting FCD patients. Future studies may investigate roles of microglia in FCD pathophysiology in greater depth, since these could represent a promising therapeutic target. For example, inhibition of one of the significantly upregulated signaling pathways in FCD, JAK-STAT3, was recently shown to have profound seizure suppressive properties in mouse models of focal epilepsy^41^.

Interrogation of epilepsy-associated pathways and gene lists strongly suggested global alterations affecting synapse formation and function in FCD. Our examination of intercellular communication revealed altered glutamate and GABA-A signaling in neurons. Given the integral role of glutamatergic and GABAergic synapses in neuronal synchronization and function, unsurprisingly dysregulation of signaling pathways that affect the normal function of glutamate and GABA in neurons, may play a role in epilepsy pathogenesis^42–45^. Consistent with our findings, a prior study also described differential expression of glutamate and GABA-A receptor subunits in dysplastic and heterotopic neurons in FCD^46^, although to our knowledge, cell type-specific changes with clear direction of intercellular communication have never been described before.

The enrichment of *MTOR* variant-carrying cells in the L2/3 IT cluster and the similar trend for the *PIK3CA* variant, can be potentially explained by three different processes: 1) FCD variants occur during later cortical neurogenesis and thus affect progenitors that preferentially produce upper-layer neurons; 2) variant-carrying cells with other cell identities may be negatively selected in terms of survival; and 3) the mutation confers a bias to the progenitor cells, causing them to produce more upper-layer neurons compared to non-mutant progenitors at a similar timepoint (Fig. 6). Although the second and third hypotheses may appear contradictory with the preserved cell type composition in cases and controls in our data (Fig. 1e), the variant-carrying cells are a minority in FCD brains and so a potential increase in the proportion of upper-layer neurons may not be detectable with our current sample size. Given the focal and localized nature of the dysplastic cortex in FCD, we speculate that FCD variants may first arise in dorsal forebrain progenitors that produce radially migrating neurons and glial cells.

**Figure 6.**
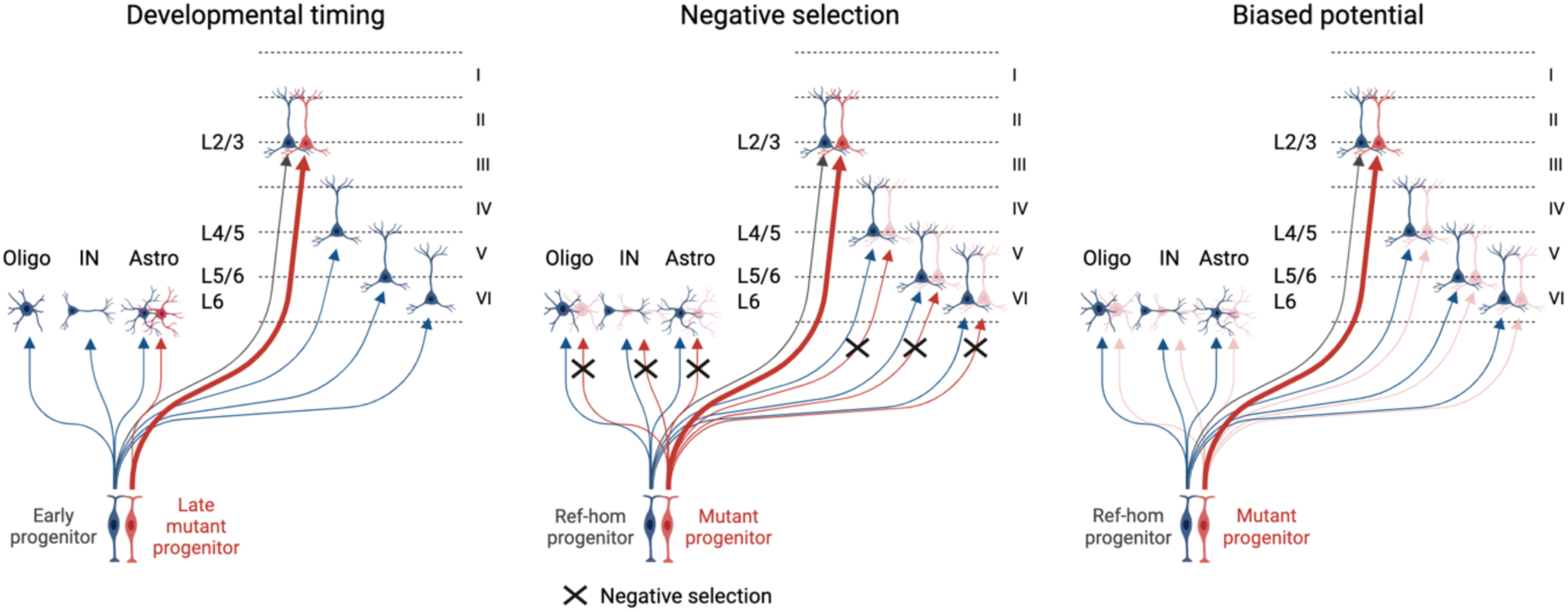
Potential mechanisms of mutant cell enrichment in L2/3 IT neurons. Schematic depicting three different hypothetical scenarios that may cause enrichment of mutant cells in L2/3 excitatory neurons. In the left panel, an early progenitor (blue) produces glial cells, inhibitory neurons, and excitatory neurons of all cortical layers, while a late progenitor carrying the mutation (red) produces preferentially L2/3 IT neurons and astrocytes. In the middle panel, a ref-hom progenitor (blue) produces all cell types, while a mutant progenitor (red) at the same developmental timepoint preferentially produces L2/3 IT neurons as a consequence of negative selection acting on other cell types. In the right panel, the mutation causes a bias in the progenitor potential, causing an increased production of L2/3 IT neurons compared to the ref-hom counterpart.

One of the limitations of our study is the low sensitivity of GO-TEN, which in its current form allows to genotype ∼21% of the sequenced cells in the high-sensitivity mode. The sensitivity may be even lower for low VAF variants. This leads to reduced specificity of downstream transcriptional analyses that aim at detecting genotype-associated gene expression changes. A second limitation is the technically demanding nature of the GO-TEN technology so that not all alleles are accessible for genotyping. Finally, our regression analysis reveals that certain cell types are preferentially genotyped for a given gene, probably due to isoform-specific expression. Future developments will aim at overcoming these limitations by increasing sensitivity and by widening the applicability spectrum.

Despite these limitations, our study highlights the importance of investigating both cell-autonomous and non-cell-autonomous mechanisms of somatic variants in mosaic FCDs, since both aspects are likely involved in disease pathophysiology. Furthermore, we illustrate the bi-directional disruption of intercellular communication in neurons that likely plays a key role in epileptogenesis and chronic epilepsy. Our single cell genotyping strategy focused on GoF variants in two well-established FCD genes, *PIK3CA* and *MTOR*, could be applied at a larger scale in future studies to address the potentially different impact of specific genes and variants in FCD and help generate more effective, targeted pharmacological treatments.

## Methods

### Patient cohort and human tissue

This study was approved by the Institutional Review Board (IRB) of Boston Children’s Hospital (05-05-76R and 09-02-0043). Subjects were identified and evaluated in a clinical setting, and post-surgical brain specimens were collected for research after obtaining written informed consent. Twenty-three total patients were included: ten with FCD2, seven with HME, one with FCD1A, one with FCD not otherwise specified (NOS), and four with mesial temporal lobe epilepsy (TLE) and unaffected temporal neocortex on pathology that were used as controls. Of the 23 patients included, 16 had a genetic diagnosis^3, 13^, including 1 germline variant in *DEPDC5*, and 15 somatic variants in *MTOR*, *AKT3*, *TSC1*, *PIK3CA* and *SLC35A2* (see Table 1 and Extended Data Table 1 for clinical details and the variants previously identified). Deidentified patient samples were also obtained from surgical resections following written informed consent from Thomas Jefferson University Hospital (IRB approval 98.0550), Cleveland Clinic Epilepsy Biospecimen Bank (IRB approval 12-1000) and Melbourne Brain Centre (Austin Health IRB approval 02961).

Neurotypical control post-mortem prefrontal cortex tissue was obtained from a 15-year-old female (UMB4638) and a 42-year-old female (UMB4643), both without history of neurologic disease, from the National Institute of Health (NIH) NeuroBioBank at the University of Maryland Brain and Tissue Bank (UMBTB). As approved by the University of Maryland IRB, research on these de-identified specimens and data was performed at Boston Children’s Hospital.

### Single-nuclei RNA-sequencing

Single-nuclei (sn)RNA-seq was performed using the 10X Genomics Chromium Next GEM Single Cell 3ʹ Reagent Kit v3.0 and v3.1. To obtain nuclei suspensions, fresh-frozen postoperative brain specimens and control post-mortem brain tissues were dissociated on ice in chilled nuclear lysis buffer (10 mM Tris-HCl, 0.32 M Sucrose, 3 mM MgAc_2_, 5 mM CaCl_2_, 0.1 mM EDTA, pH 8, 1 mM DTT, 0.1% Triton X-100 and 0.2 U/μl RNase inhibitor) using a dounce homogenizer. Homogenates were layered on top of a sucrose cushion buffer (1.8 M Sucrose, 3 mM MgAc_2_, 10 mM Tris-HCl pH 8, 1 mM DTT) and ultra-centrifuged for 1 hour at 30,000 rcf. Pellets containing nuclei were resuspended in 500 μl ice-cold 1X PBS supplemented with 3 mM MgCl_2_ and 0.2 U/μl RNase inhibitor, then filtered through a 40 μm cell strainer. Nuclei suspensions were centrifuged at 500 rcf and 4 °C for 5 minutes, and pellets re-suspended in 80 μl nuclease-free H_2_O. For some preparations, an intermediate step consisting of DAPI staining and fluorescence-activated nuclear sorting (FANS) was performed before centrifuging for 5 minutes at 500rcf. Nuclei sorting was performed on a FACS Aria II cell sorter equipped with BD FACSDiva software, selecting all DAPI-positive nuclei (SSC-A and FSC-A gate boundaries were set at 10^3^ and 30, respectively). Around 30,000 nuclei were used to load the 10X Chromium, and were processed the same day for gel-bead in emulsion (GEM) generation, barcoding, and cDNA amplification, following manufacturer instructions. Libraries were sequenced (paired-end single indexing) on an Illumina NovaSeq6000 targeting ∼50,000 read-pairs per single-nucleus.

### Single-nucleus RNA-sequencing gene expression quantification, quality control, and processing

Alignment and gene expression quantification of snRNA-seq data was performed with the 10X CellRanger pipeline (v8.0.1) based on the GRCh38 reference genome using all default parameters and with introns included (--include-introns flag). As previously reported^17^, ambient RNA contamination can confound analysis of snRNA-seq data. To address this issue, we utilized CellBender (v0.3.0)^47^ with all default parameters to adjust raw counts based on automatic detection of likely contaminant reads. We additionally used scrublet (v0.2.3)^48^ to detect and remove likely doublets. Seurat (v4.4.0)^49^ was then used for all subsequent downstream analyses. In brief, high-quality nuclei were extracted based on the following criteria: 1) percent mitochondrial gene expression < 5%; 2) percent ribosomal gene expression < 5%; and 3) number of expressed genes > 300 (Extended Data Fig. 1b). snRNA-seq was then normalized (*NormalizeData)* and scaled (*ScaleData)*, with regression of the covariates of percent mitochondrial gene expression, percent ribosomal gene expression, number of expressed genes (nFeature_RNA), and number of transcripts (nCount_RNA). All samples were integrated using Harmony (v1.2.0)^50^ (Extended Data Fig. 2) and unsupervised clustering was performed using Louvain’s algorithm (*FindClusters)*.

DropletQC (v0.0.0.9000)^51^ was used to calculate the intronic ratio per-nuclei and unsupervised clusters with a mean intronic ratio < 0.4 were filtered to further eliminate ambient contamination and damaged droplets. Using canonical brain cell type marker genes^18^, cell-type annotation was performed for each high-quality, retained unsupervised cluster.

### Differential cell type composition analysis

Cell type composition for each sample was calculated independently. To test whether any cell types show differential composition among controls, FCD1 cases, and HME/FCD2 cases, a one-way ANOVA test was performed in R using the *aov* function from the R package *stats* (v4.3.1) and the following formula for each cell type: cell type proportion per sample ∼ disease status (i.e. control, FCD1, HME/FCD2). A Benjamni-Hochberg correction was used to control for multiple hypothesis testing.

### Differential gene expression and pathway enrichment analysis

Differential expression analysis was performed using Seurat (v4.4.0)’s FindMarkers function (wilcox test). Only genes expressed in at least 10% of nuclei within a particular comparison were considered (min.pct = 0.10) and no log2 fold-change threshold was imposed (logfc.threshold = 0). Default parameters were used otherwise. To understand pathways that were differentially expressed among comparisons of interest, we utilized gene set enrichment analysis as implemented in the R package fgsea (v1.28.0), with genes ranked in descending order by average log2 fold-change. A Benjamjni-Hochberg correction was used to control for multiple hypothesis testing. Relevant ontologies were obtained as follows:

1) Hallmark pathways: MSigDB (https://www.gsea-msigdb.org/gsea/msigdb)
2) Genes associated with epilepsy: DisGenNet (https://www.disgenet.com/) and Macnee et al., 2023^25^
3) SynGO pathways: SynGO (https://www.syngoportal.org/)
4) Autism genes: SFARI Gene (https://gene.sfari.org/)
5) Microglial state marker genes: Friedman et al., 2018^22^.

### GO-TEN single nucleus genotyping

Two FCD (FC5501 and FC5801) and one HME (E174) cases were used for GO-TEN analysis. Excess full-length cDNA from 10X Genomics Next GEM Single Cell 3′ GEM Kit v3.1 workflow was used as input for GO-TEN genotyping. 10 ng of excess cDNA after the amplification step was used to perform hemi-nested long-range PCR to capture the target variant as well as the unique 10X barcode and UMI. The PrimeSTAR GXL DNA Polymerase was used, following the manufacturer’s manual, given optimal performance at the desired amplicon length. The long-range PCR product was then used for long-read sequencing library preparation using the ONT Kit12 chemistry. The final library was sequenced on one MinION Flow Cell.

Genotyping of known variant sites was performed as follows. MinION reads were aligned with MiniMap2 (v2.24) to the GRCh38 human reference genome. Each read was parsed based on the expected structure of the read (i.e. known primer sequences, 10X cell barcodes and UMIs, and poly-A tail reads) to identify the allele present at the known variant site. A small portion of reads were identified as containing neither the reference or expected variant allele; these reads were discarded as they likely represent sequencing errors. Reads that did not contain a 10X CBC that was present in the final processed snRNA-seq dataset were also discarded. To optimize for stringency of genotyping while still maximizing statistical power to detect biological differences between wild-type and mutant nuclei, we then implemented two downstream genotyping approaches. In the “strict” high-specificity genotyping approach, we further filtered reads that did not contain a 10X UMI identified in our snRNA-seq data as corresponding to the gene we were attempting to genotype (i.e. *PIK3CA* or *MTOR*). Such an approach minimizes the chance of genotyping reads that are off-target or have an unexpected structure (although the vast majority of these reads will be filtered by our additional, aforementioned filtering criteria) but also may discard many on-target reads whose associated 10X UMI was filtered from our snRNA-seq data despite the overall 10X CBC it corresponds to not being filtered during our snRNA-seq data processing. In our “loose” high-sensitivity genotyping approach, we did not apply the latter additional filtering strategy. Finally, we used the Bayesian genotyper derived from MosaicHunter^35, 36^ in two steps: 1] determine the UMI-level genotype (reference or mutant) by aggregating multiple reads with the same UMI, and 2] determine the nuclei-level genotype (reference-homozygous or heterozygous mutant) by considering multiple UMIs within the same nuclei. In each step, the base quality and read/UMI count were considered as the likelihood in the Bayesian model, while the mutant cell fraction estimated from bulk DNA sequencing were additionally considered as the prior probability in the second step.

To identify cell types carrying our variants of interest (*variant cell types)*, we only utilized strict genotyping calls to maximize our confidence in identifying the developmental origin of these mosaic variants. When performing differential expression analysis, we only considered variant cell types identified based on our strict genotyping approach. To maximize our statistical power to identify differentially expressed genes and pathways between mutant and wild-type cells of these variant cell types, we utilized all nuclei of these variant cell types that were confidently genotyped as either reference-homozygous or heterozygous in our high-sensitivity genotyping approach.

### Genotyping efficiency analysis

To identify covariates associated with genotyping efficiency (i.e. the probability a nucleus was genotyped confidently, regardless of whether it was identified as reference-homozygous or heterozygous mutant), we fit the following logistic regression model (R *stats* package v4.3.1 *glm* function) across all nuclei successfully genotyped based on both our genotyping approaches (Extended Data Fig. 4a), as described above: nucleusWasGenotyped (binary outcome; 1 = was genotyped and 0 = was not genotyped) ∼ expression of target gene + cell type.

## Code availability

Scripts are available upon reasonable request.

## Data availability

Previously generated^14^ de-identified human single-nucleus RNA-seq data have been deposited at the NIAGADS DSS under the accession number NG00162. They are available upon request if access is granted. To request access, contact the NIGADS DSS (https://dss.niagads.org/). Newly generated de-identified human single-nucleus RNA-seq data are being deposited at dbGaP and will be available upon request if access is granted at https://dbgap.ncbi.nlm.nih.gov/aa/.

## Author contributions

S.B., E.A.S, M.T. and S.K. conceived the study. S.B. and S.K. established the patient cohort, collected surgical specimens and performed snRNA-sequencing experiments. E.A.S. conceived the GO-TEN protocol, performed GO-TEN experiments, and contributed to GO-TEN bioinformatic analyses. B.H.C. helped with GO-TEN experiments. Z.Z. helped troubleshoot GO-TEN experiments. M.T. led and performed all bioinformatic analyses, including snRNA-seq and GO-TEN. Q.H., A.Y.H. and S.K. contributed to bioinformatic analyses. T.E.G., and M.S.H. performed molecular data acquisition and analyses for one sample in the study cohort. D.C.R., and S.A.M. performed clinical phenotyping for one case in the study cohort. A.H.P. and R.J.B. provided a subset of epilepsy surgical resections in the study cohort. A.H.P, A.M.D., and E.Y. performed phenotypic analysis and summary of clinical data. S.B., M.T. and S.K. interpreted data analysis and generated manuscript figures. S.B. wrote the manuscript, greatly helped by M.T., S.K. and C.A.W. All authors edited the manuscript. S.K. and C.A.W. directed the research, with the help of S.B.

### Acknowledgments

We thank the patients and their families for their invaluable donations to the advancement of science. We thank J.E. Neil, the Boston Children’s Hospital Core Repository for Neurological Disorders, and the NIH Neurobiobank at the University of Maryland Brain and Tissue Bank for facilitating tissue access. We thank Prof. Brenda E. Porter (Department of Neurology, Stanford University School of Medicine), Prof. Dr. Imad Najm, Prof. Dr. Zhong Ying, Dr. Robyn Busch, Dr. Ingmar Blümcke and the Cleveland Clinic Epilepsy Biospecimen Bank, Thomas Jefferson University, and the Melbourne Brain Centre at Austin Health for providing epilepsy surgical resections included in the study cohort. We thank the Boston Children’s Hospital IDDRC Molecular Genetics Core Facility, supported by NIH award U54HD090255 from the National Institute of Child Health and Human Development. Funding: supported by R01AG070921 from the NIA and R01NS032457 from the NINDS to C.A.W. and R01CA269805, R01HG012573, and UM1DA058230. C.A.W. is an Investigator of the Howard Hughes Medical Institute. S.B. was supported by the Manton Center for Orphan Disease Research at Boston Children’s Hospital, and was supported by the Horizon2020 Research and Innovation Program Marie Skłodowska-Curie Actions (MSCA) Individual Fellowship (grant agreement no. 101026484—CODICES). S.K. was supported by NINDS award K08NS128272, Physician Scientist Fellowship from the Doris Duke Charitable Foundation, and Career Award for Medical Scientists from the Burroughs Wellcome Fund. M.T. and E.A.S. were supported by National Institutes of Health grants T32GM007753 and T32GM144273. Z.Z. was supported by PRMRP Discovery Award W81XWH2010028, the Edward R. and Anne G. Lefler Center postdoctoral fellowship and the American Heart Association Career Development Award 23CDA1046074. A.Y.H was supported by R56AG079857 from the NIA and Alzheimer’s Association Research Fellowship. This study was funded by the National Health and Medical Research Council Australia (NHMRC). M.S.H. was supported by an NHMRC Ideas Grant (2012287) and NHMRC Project Grants (1129054 & 1079058). We thank Dr. Michael R Sperling and Dr. Thomas N. Ferraro for their assistance in acquiring the TLE patient samples from Thomas Jefferson University Hospital and acknowledge NINDS funding R01NS49306 to RJB supporting this effort.

## Extended Data

**Extended Data Table 1.** Additional clinical details related to the patient cohort included in the study.

**Extended Data Figure 1.**
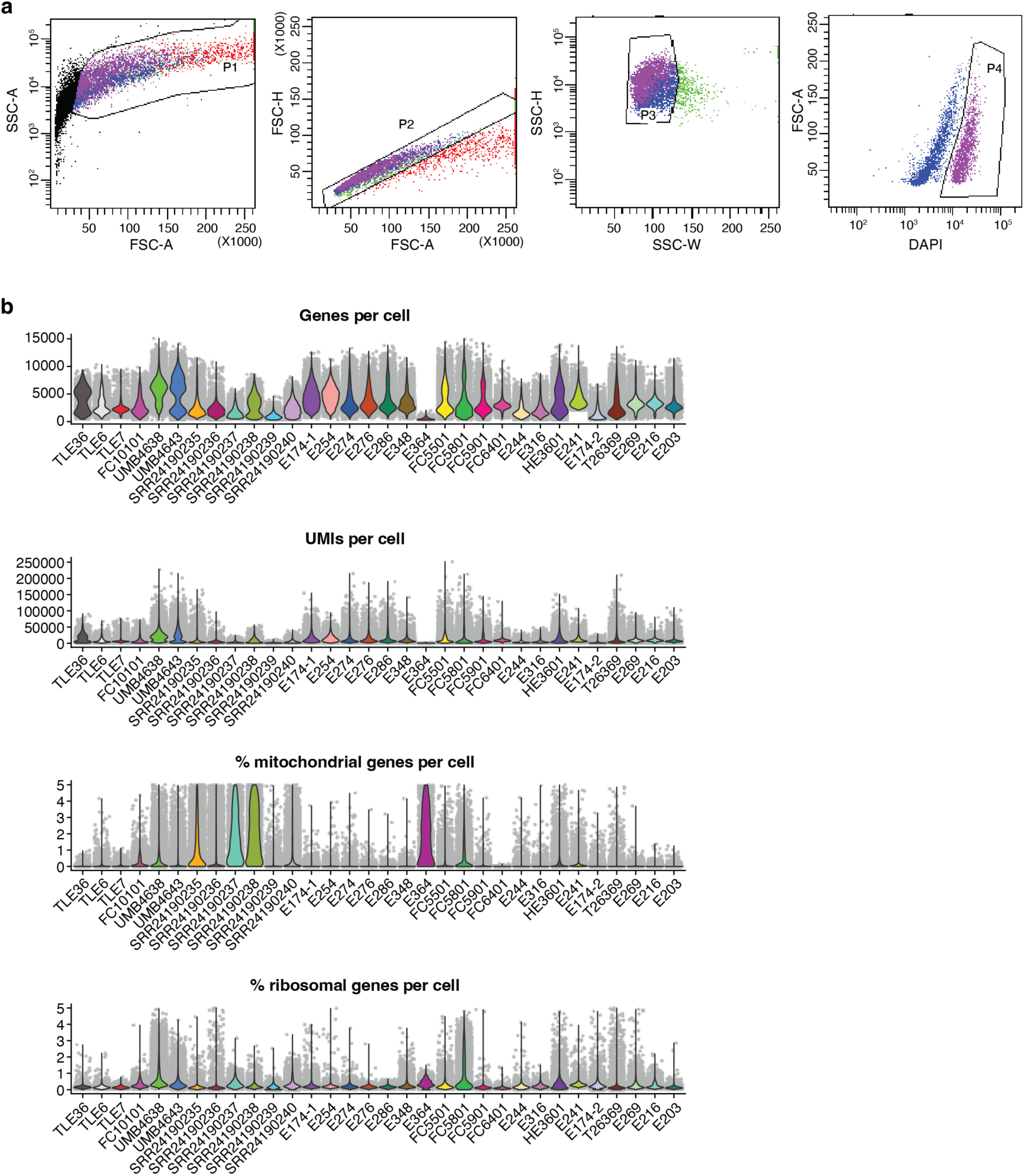
DAPI sorting and single-nucleus RNA-sequencing data quality. (**a**) Representative images of the gating strategy performed for DAPI fluorescent activated nuclear sorting upstream of single-nuclei RNA-sequencing. (**b**) Distribution of snRNA-seq quality parameters for each sample included in the study.

**Extended Data Figure 2.**
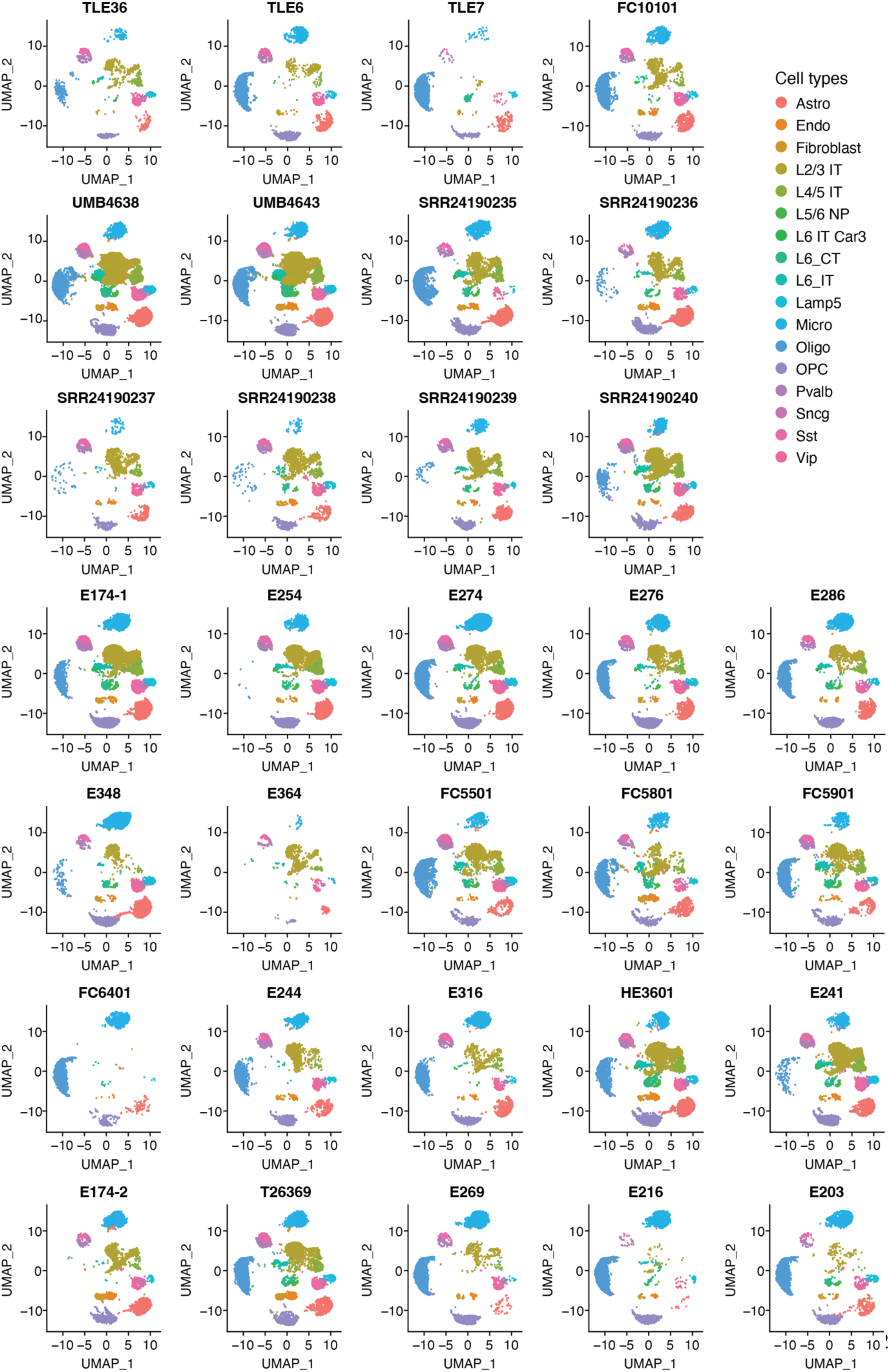
Single-nucleus RNA-sequencing data integration. Uniform Manifold Approximation and Projection (UMAP) dimension reductions are reported for each individual sample. Data integration was performed with Harmony (v1.2.0).

**Extended Data Figure 3.**
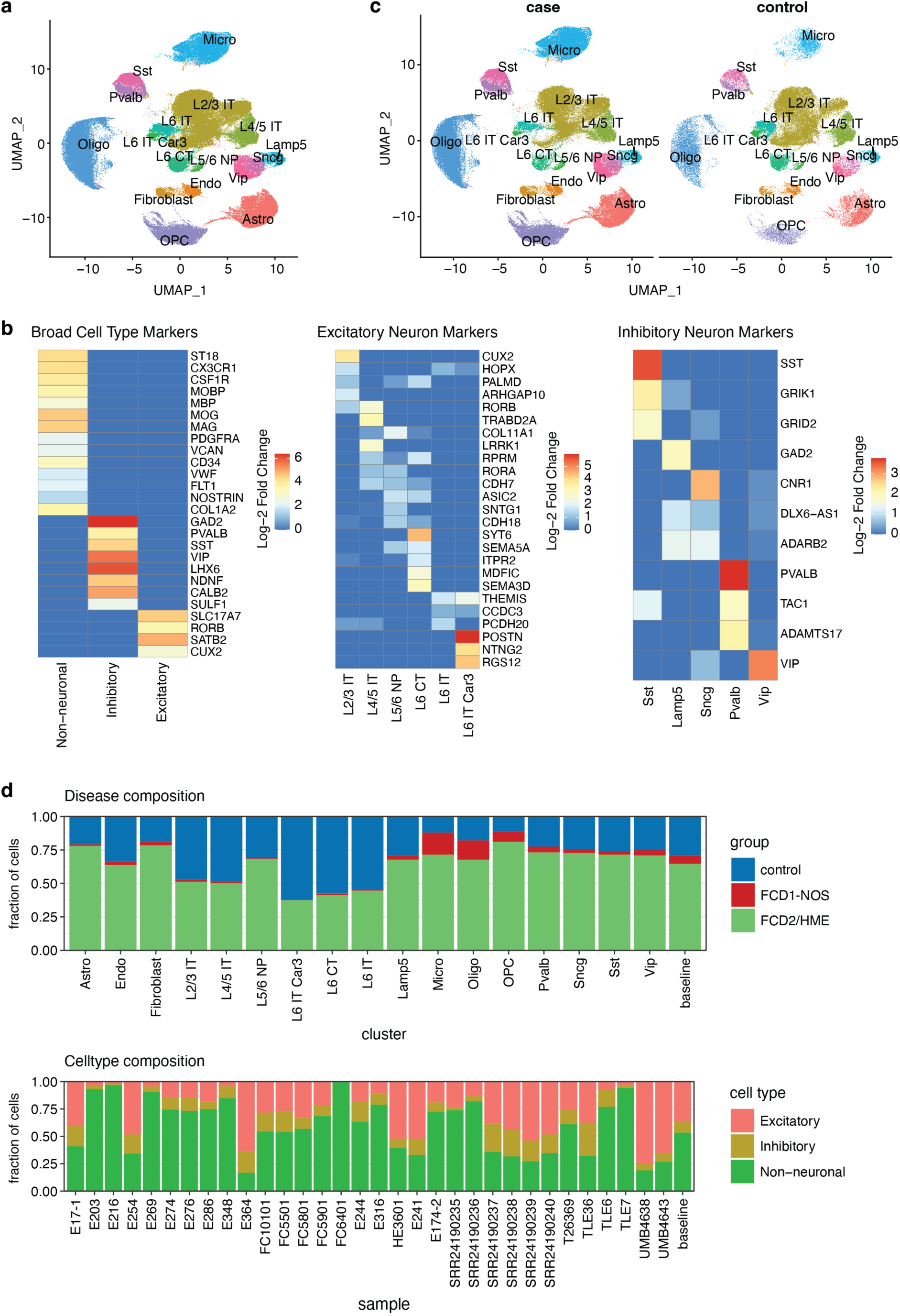
Analysis of cell types including data from Chung et al. (**a**) UMAP showing cell type clusters identified in the integrated disease and control samples. (**b**) Heatmaps showing the expression of known cell type markers used for annotation. (**c**) UMAPs showing conserved cell type clusters between cases and controls. (**d**) Contribution of each disease and control group to cell types (top panel) and the contribution of each broad cell type group to samples (bottom panel). Differential cell type composition analysis was done using a one-way ANOVA test, which showed non-significant shifts.

**Extended Data Figure 4.**
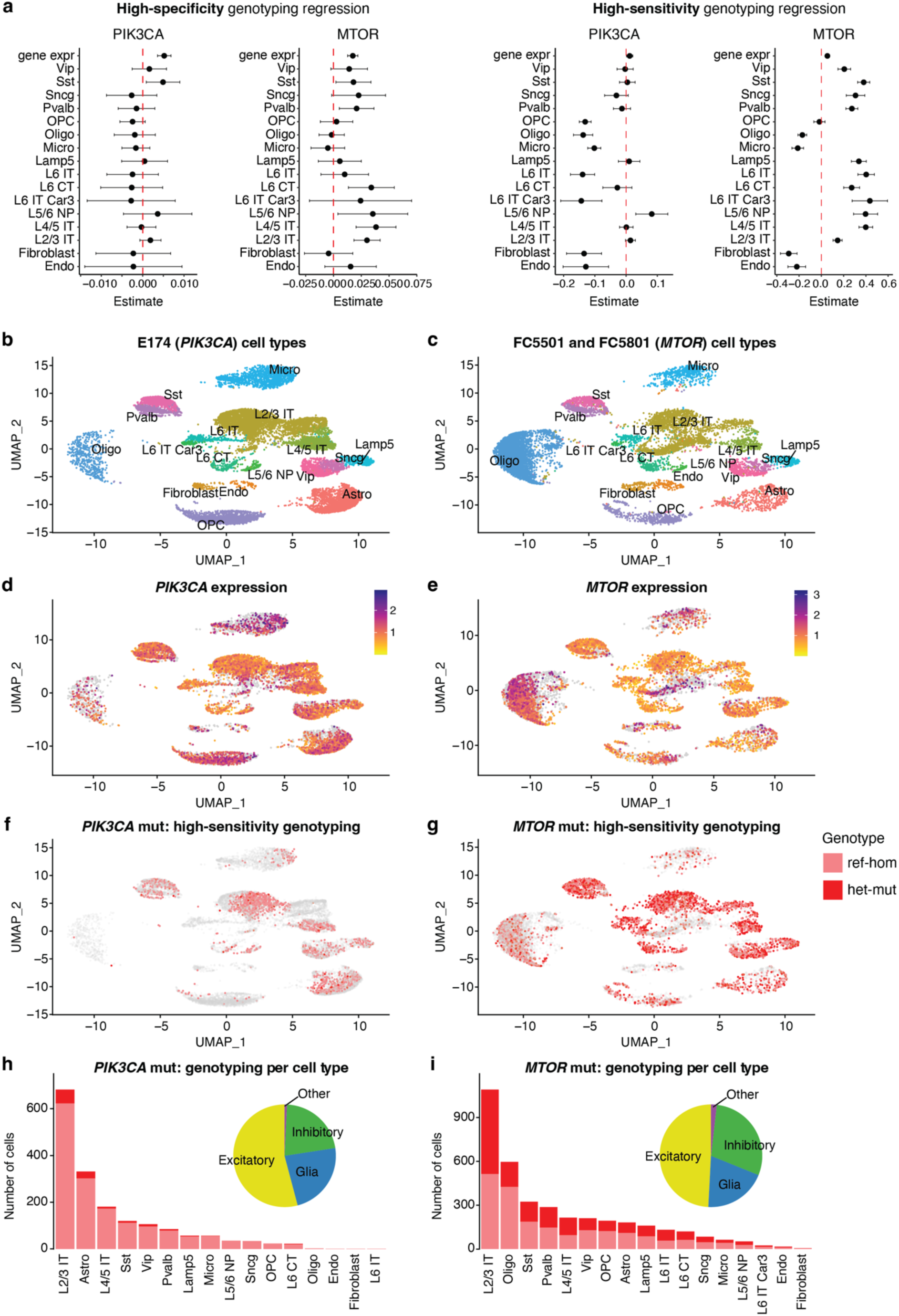
High-sensitivity GO-TEN genotyping. (**a**) Logistic regression estimates for covariates tested for association with GO-TEN genotyping efficiency using the high-specificity and high-sensitivity approaches. (**b-c**) UMAPs showing cell type clusters for cases genotyped with GO-TEN. (**d-e**) *PIK3CA* and *MTOR* expression across cell types based on snRNA-seq. (**f-g**) UMAPs showing *PIK3CA* and *MTOR* reference homozygous and heterozygous mutant cells genotyped with the GO-TEN high-sensitivity approach. (**h**-**i**) Number of cells genotyped with the GO-TEN high-sensitivity approach as reference homozygous and heterozygous mutant. Pie charts show the contribution of broad cell categories to mutant cells.

